# Pleiotropic enhancers are ubiquitous regulatory elements in the human genome

**DOI:** 10.1101/2022.01.25.477532

**Authors:** Ian Laiker, Nicolas Frankel

## Abstract

Enhancers are regulatory elements of genomes that determine spatio-temporal patterns of gene expression. The human genome contains a vast number of enhancers, which largely outnumber protein-coding genes. Classically, enhancers have been regarded as highly tissue-specific. However, recent evidence suggests that many enhancers are pleiotropic, with activity in multiple developmental contexts. Yet, the extent and impact of pleiotropy remain largely unexplored. In this study we predicted active enhancers across human organs based on the analysis of both eRNA transcription (FANTOM5 consortium datasets) and chromatin architecture (ENCODE consortium datasets). We show that pleiotropic enhancers are pervasive in the human genome and that most enhancers active in a particular organ are also active in other organs. In addition, our analysis suggests that the proportion of context-specific enhancers of a given organ is explained, at least in part, by the proportion of context-specific genes in that same organ. The notion that such a high proportion of human enhancers can be pleiotropic suggests that small regions of regulatory DNA contain abundant regulatory information and that these regions evolve under important evolutionary constraints.

**Significance statement:** The human genome contains a vast number of regulatory elements, named enhancers, that control the tempo and mode of gene expression. Classically, enhancers have been regarded as genetic elements that are active in a single organ, but recent evidence suggests that many enhancers in animal genomes are active in multiple organs (i.e., are pleiotropic). Here we shed light on the architecture of non-coding human DNA by showing that a large percentage of the enhancers of the human genome are pleiotropic and that the majority of enhancers active in a particular organ are pleiotropic. This suggests that small regions of regulatory DNA may contain abundant regulatory information and that these regions evolve under important evolutionary constraints.

## Introduction

Enhancers are regulatory elements of the genome, in general smaller than 1 kb, that determine spatio-temporal patterns of gene expression (Levine 2010). Enhancers are activated and repressed by the binding of transcription factors (TFs), which recognize specific motifs in the DNA named transcription factor binding sites (TFBSs)(Hill et al. 2021). Upon activation, enhancers interact with the basal transcriptional machinery at the core promoter, boosting mRNA synthesis of the target gene(s)(Panigrahi & O’Malley 2021). There are several features that distinguish active enhancers: (i) it has been shown that nucleosomes flanking active enhancers usually contain an acetyl group in lysine 27 of histone H3 (H3K27ac) (Creyghton et al. 2010) and a single methyl group in lysine 4 of histone H3 as well (H3K4me1)(Rada-Iglesias et al. 2011), which often coincides with the absence of the trimethylated form of H3 lysine 4 (H3K4me3)(Lidschreiber et al. 2021), (ii) it has been demonstrated that enhancer DNA may be transcribed bidirectionally, generating short capped non-polyadenylated RNAs (‘eRNAs’)(Sartorelli & Lauberth 2020), and (iii) it is typically observed that active enhancers are depleted of positioned nucleosomes, thus having a conformation that is known as ‘open chromatin’ (Klemm et al. 2019). Together, these features can be used to identify enhancers genome-wide (Kimura 2013; Kristjánsdóttir et al. 2020; Minnoye et al. 2021).

The human genome contains a vast number of enhancers, which largely outnumber protein-coding genes (Pennacchio et al. 2013; Heintzman et al. 2009). It has been shown that a large percentage of SNPs associated with human diseases fall within regions that are predicted to be enhancers (Corradin & Scacheri 2014), and, in the same vein, numerous cases in which mutations in enhancers cause disease have been reported (Smith & Shilatifard 2014). Although thousands of non-coding regions have been identified as putative enhancers, very little is known about the function of most of these predicted enhancers (Gasperini et al. 2020). Indeed, the complexity of enhancer function in the human genome is just beginning to be comprehended. For example, recent studies uncovered that enhancers usually regulate multiple genes and that many enhancers bypass their nearest core-promoters to interact with distant core-promoters (Reilly et al. 2021; Nasser et al. 2021). The fact that regulatory information within enhancer DNA can be dense and pleiotropic (Fuqua et al. 2020; Kvon et al. 2020; Xin et al. 2020; Le Poul et al. 2020) also illustrates regulatory complexity.

The classic conception in which enhancers have strict tissue-specific activities (i.e., modular or context-specific activities) has been called into question by recent experimental data (Fuqua et al. 2020; Infante et al. 2015; Lewis et al. 2019; Lonfat et al. 2014; Preger-Ben Noon et al. 2018; Xin et al. 2020). Current evidence suggests that a large proportion of enhancers in vertebrate genomes are active in multiple developmental contexts (Kittelmann et al. 2021; Sabarís et al. 2019), which probably means that a significant number of enhancers in the human genome are pleiotropic (i.e., have roles in multiple organs and/or developmental stages). Although evidence suggest that pleiotropy in regulatory DNA is pervasive, the extent of enhancer pleiotropy in the human genome remains largely unexplored. In this sense, it would be useful to have a reliable estimate of the percentage of pleiotropic enhancers in the human genome and to explore the landscape of pleiotropy in each organ.

To assess the magnitude of pleiotropy in the regulatory genome, in this study we estimated the extent of enhancer reuse and analyzed the abundance of pleiotropic enhancers per organ. We show that pleiotropic enhancers are pervasive in the human genome and that the majority of enhancers active in a particular organ are also active in other organs. In addition, our analysis suggests that the proportion of context-specific enhancers of a given organ is explained, at least in part, by the proportion of context-specific genes in that same organ.

## Results

### A large proportion of predicted human enhancers are pleiotropic

We first set out to estimate the number of pleiotropic enhancers in the human genome by using information from the FANTOM5 (Andersson et al. 2014) and ENCODE (Dunham et al. 2012) projects. We grouped ENCODE and FANTOM biosamples into organs using UBERON anatomy ontology (Mungall et al. 2012) (see tables S1 and S2). For FANTOM5 data, putative enhancers were defined as DNA regions transcribed bidirectionally above a certain threshold in at least one of 41 organs (see table S1 for a list of samples and organs). For ENCODE data, we defined putative enhancers as DNA regions with open chromatin that contain the epigenetic mark H3K27ac in at least one of 20 organs (see table S2 for a list of samples and organs). Although ENCODE and FANTOM5 predictions are derived from different biosamples and disparate genomic analyses, 40-60% of the FANTOM5 enhancers are contained within the larger ENCODE enhancer set in 16 common organs (testis is an exception, see Figure S1). Also, we compared ENCODE enhancers with the ‘ABC enhancer set’, which was defined based on chromatin accessibility, the presence of the epigenetic mark H3K27ac and the interaction of the DNA region with core promoters (Fulco et al. 2019). We determined that, for 14 common organs, the overlap between our enhancer set predicted with ENCODE data and ABC enhancers ranges between 43% and 83% (Figure S1). Altogether, comparisons between different enhancer sets suggest that most of our enhancer predictions based on ENCODE data are robust.

By intersecting genomic information from different organs we aimed to estimate the proportion of pleiotropic enhancers in the human genome. Hence, we considered that a pleiotropic FANTOM5 enhancer is a DNA region transcribed bidirectionally in two or more organs. For ENCODE data, we delimited consensus elements considering the different organs in which an enhancer was predicted to be active, based on the presence of open chromatin and the H3K27ac mark (see details of the pipeline in figure S2 and Materials and Methods, and figure S3 for the size distribution of consensus elements). In this case, we define a pleiotropic enhancer as a consensus element active in two or more organs. To evaluate how the proportion of pleiotropic enhancers changes with the number of organs considered we took combinations of organs from 2 to *n* out of the pool of ENCODE organs (*n*=20) and FANTOM5 organs (*n*=41) and calculated the proportion of enhancers active in 2 or more organs. This approach allows us to comprehend how the proportion of pleiotropic enhancers changes depending on the number of organs considered and, at the same time, to validate our methods. We observed that the median of predicted pleiotropic enhancers increases monotonically with the number of intersected organs, both for FANTOM5 and ENCODE data (Figure 1). Although enhancer sets predicted with ENCODE and FANTOM5 data are different (∼50% of the FANTOM5 enhancer set is contained within the larger ENCODE enhancer set and, as well, the FANTOM5 enhancer set represents enhancers active in more organs than the ENCODE set) both the shape and the asymptote of the two hypothetical curves are remarkably similar (Figure 1). Furthermore, the implication that more than 40% of the predicted enhancers in the human genome are pleiotropic (active in at least two organs) is noteworthy as well (Figure 1).

**Figure 1.**
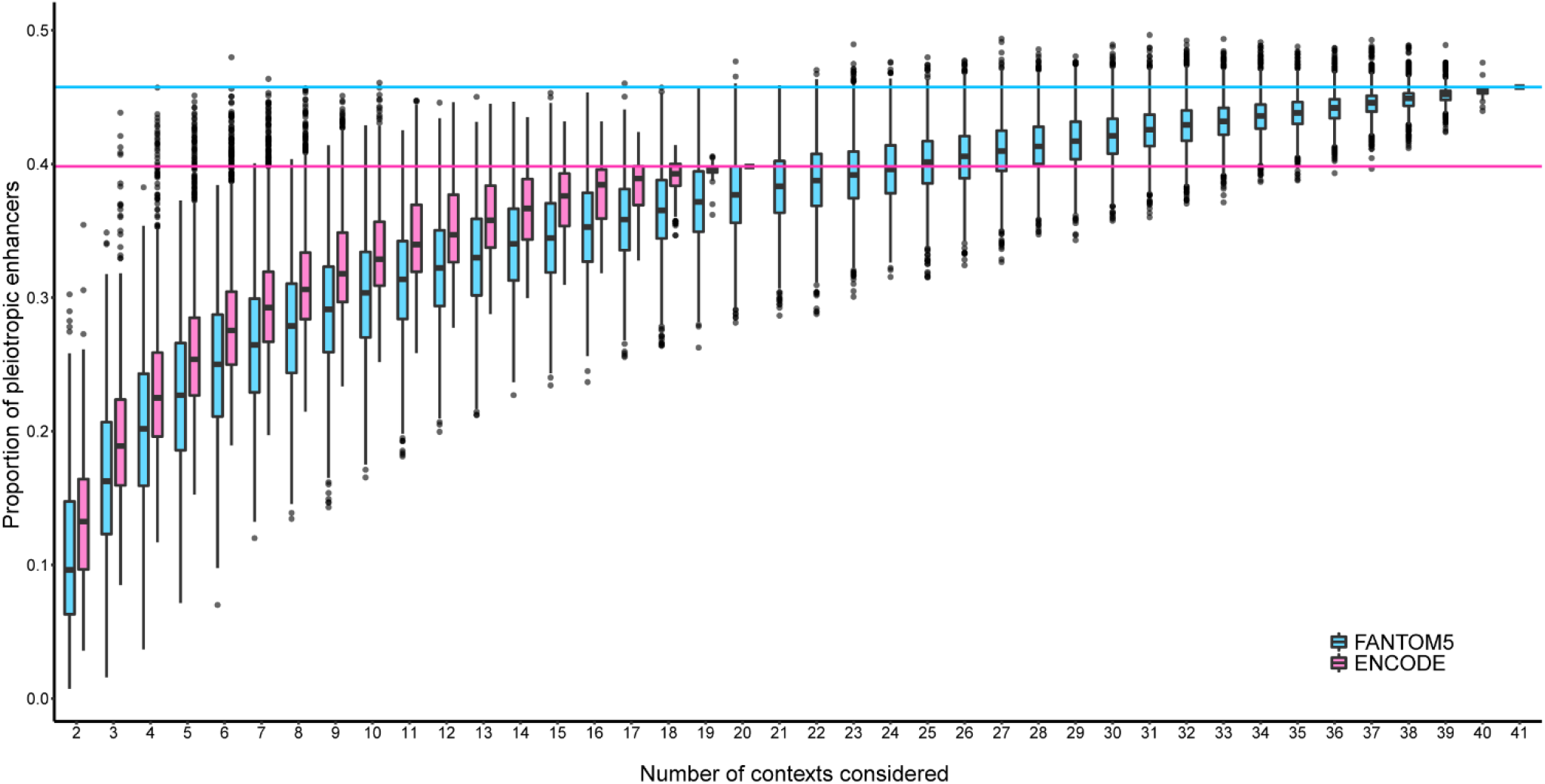
A large proportion of predicted human enhancers are pleiotropic. Proportion of pleiotropic enhancers as a function of the number of contexts considered in FANTOM5 (cyan) and ENCODE (magenta) enhancer sets. Boxplots indicate the median (horizontal line), interquartile range (box), observations within ± 1.5xinterquartile range (whiskers) and outliers (dots). The proportion of pleiotropic enhancers when considering all organs in a single comparison are marked with a magenta line (ENCODE) and a cyan line (FANTOM5).

Given that organs are comprised of different cell types and that some cell types are present in multiple organs (e.g., fibroblasts and endothelial cells), the high proportion of pleiotropic enhancers calculated with bulk organ data could be a mere consequence of considering enhancers that are active in cell types common to many organs. To rule out this possibility, we analyzed enhancers identified in primary cell cultures by the FANTOM5 project (Table S1) (Andersson et al. 2014), combining the same cell types from different organs as a single context (see Table S1). Notably, when considering these 39 distinct cell types, the percentage of pleiotropic enhancers turns out to be 68%. We performed the same analysis (Table S2) using distal enhancers (cCRE-dELS) defined by the ENCODE consortium in 17 cell types (Moore et al. 2020). When considering all 17 cell types, the percentage of pleiotropic enhancers is 39%. In order to determine whether results from cells and organs are comparable, we contrasted the degree of pleiotropy of enhancers obtained with FANTOM5 cell types data with that calculated using FANTOM5 organ data (Figure S4). We observed that the degree of pleiotropy of enhancers estimated with cell types data and organ data is remarkably similar (Figure S4). Altogether, these data suggest that enhancer pleiotropy is mostly due to enhancers being active in different cell types of organs, rather than enhancers being active in the same cell type present in multiple organs.

### Most enhancers active in a particular organ are pleiotropic

Next, we sought to explore the abundance of pleiotropic enhancers in the different organs. To this end, we calculated the proportion of predicted pleiotropic enhancers per organ in the FANTOM5 and ENCODE data sets (Figure 2). We observed values ranging from 43.6% to 98.1% in FANTOM5 organs (Figure 2A, table S3) and from 53.4% to 96.8% in ENCODE organs (Figure 2B, table S3). These data suggest that most enhancers active in a given organ are also active in one or more other organs. Indeed, most of the pleiotropic enhancers active in a given organ are active in more than two organs (Figure 2).

**Figure 2.**
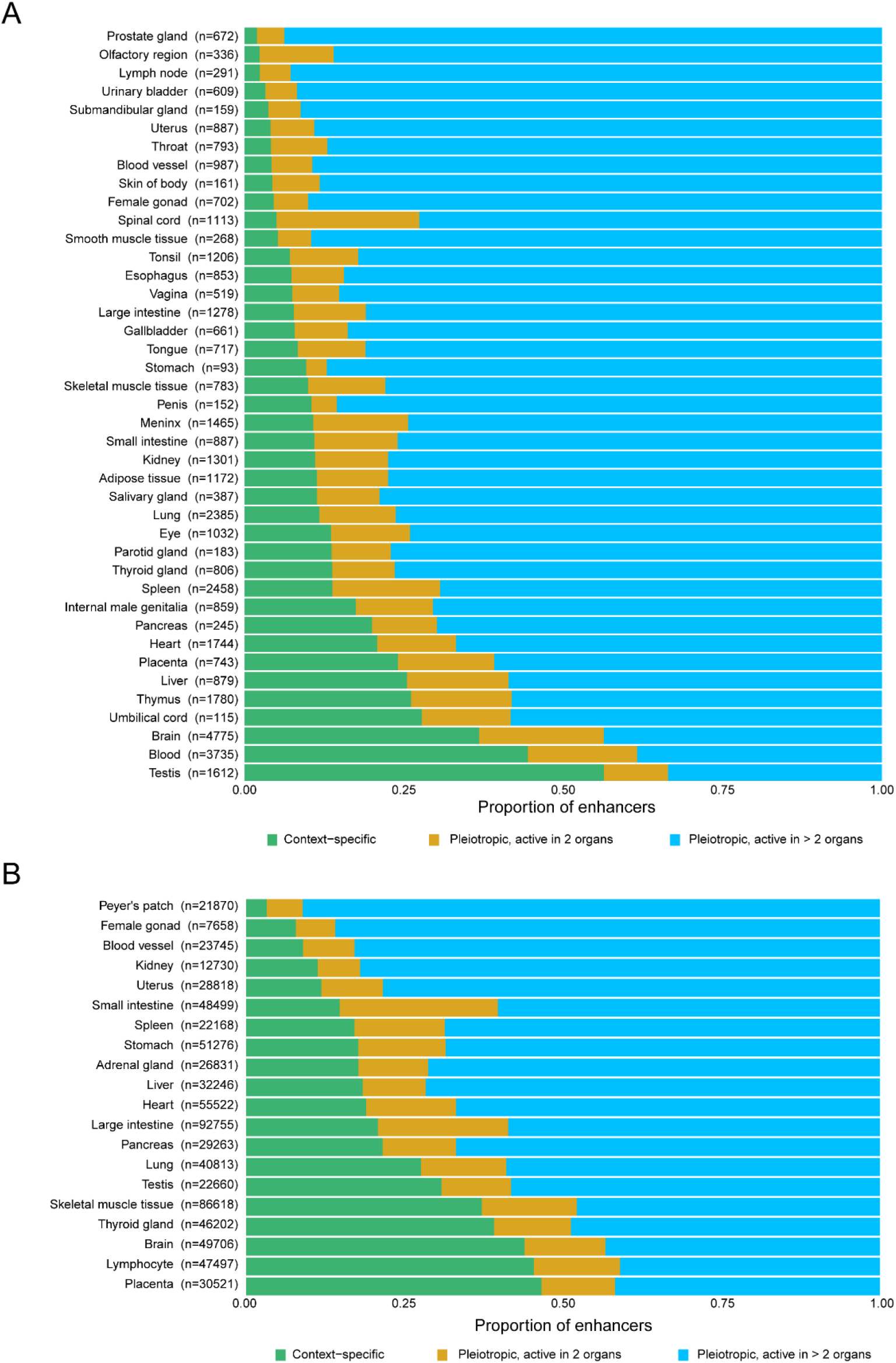
The majority of active enhancers in a particular organ are pleiotropic. Proportion of context-specific enhancers (green), pleiotropic enhancers active in 2 organs (orange) and pleiotropic enhancers active in more than 2 organs (blue) in FANTOM5 (A) and ENCODE (B) organs. The number of predicted enhancers per organ is indicated beside the name of each organ.

Given that 17 organs are shared between FANTOM5 and ENCODE datasets, we compared the estimations of the proportion of predicted pleiotropic enhancers for both analyses. We observed that they are notably concordant, especially considering that biosamples are derived from different parts of the organ and different individuals, and that enhancers were predicted through distinct methods (see Table S3). For most comparisons the difference in the estimation of the proportion of pleiotropic enhancers is smaller than 10%. However, testis is an outlier, with a difference of more than 20%. Despite differences in the estimation of the proportion of pleiotropic enhancers for testis, both FANTOM5 and ENCODE data indicate that this organ is rich in context-specific enhancers (Figure 2). Since it has been reported that testis is the organ with the most context-specific genes (Djureinovic et al. 2014; Pineau et al. 2019; Zhu et al. 2016), we reasoned that a possible cause for the high number of context-specific enhancers in the testis could be the high number of testis-specific genes and, moreover, that there could be a relationship between the number of context-specific genes and the abundance of context-specific enhancers per organ. To test this hypothesis, we computed the number of organ-specific genes in ENCODE and FANTOM5 organs (with RNA-seq and transcribed TSSs data respectively, see Materials and Methods for details) and, subsequently, we looked for a possible correlation between the proportion of organ-specific genes and the proportion of organ-specific enhancers. Remarkably, for both FANTOM5 and ENCODE organs, we found a positive correlation (Pearson’s r=0.653, p<0.001 and Pearson’s r=0.647, p<0.001, respectively) between the proportion of context-specific genes and the proportion of context-specific enhancers (Figure 3). A positive correlation is also observed when using data from FANTOM5 cell types (Pearson’s r=0.71, p<0.001).

**Figure 3.**
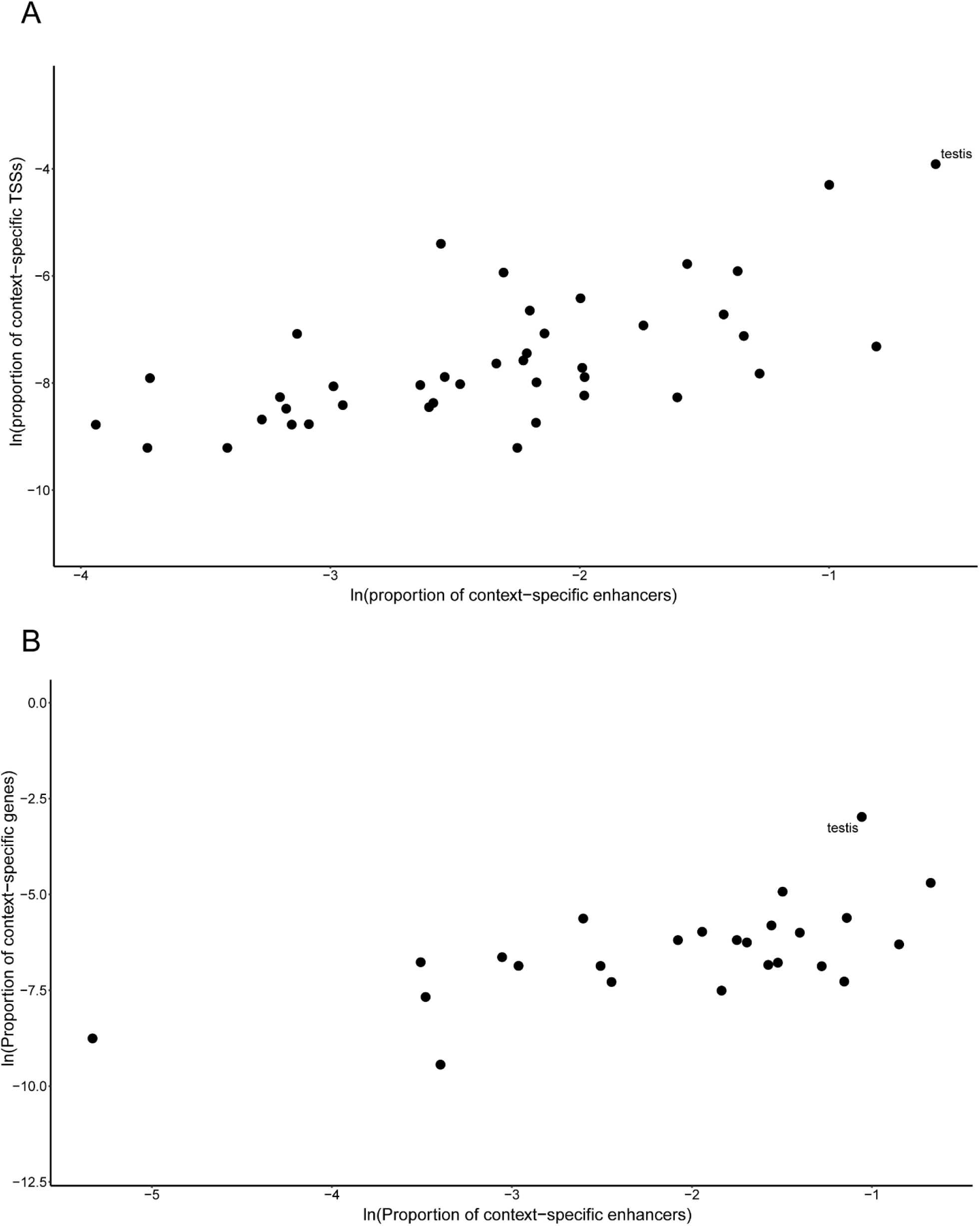
The proportion of context-specific enhancers is positively correlated with the proportion of context-specific genes. (A) Relationship between the proportion of context-specific enhancers and context-specific TSSs for the FANTOM5 dataset. Active TSSs were defined in each organ by comparing transcription levels against genomic background (see Materials and Methods for details). (B) Relationship between the proportion of context-specific enhancers and context-specific genes for the ENCODE dataset. An expression threshold was used to define expressed genes in each organ (see Materials and Methods for details).

### Pleiotropic enhancers are enriched in SNPs with high regulatory potential

Bearing in mind the fact that pleiotropic enhancers have more density of TF binding motifs (Fish et al. 2017; Singh & Yi 2021), we reasoned that a way to provide a functional validation for the pleiotropic function of enhancers was to analyze a possible connection between the degree of pleiotropy of enhancers and the abundance of genetic variants related to gene regulation. Thus, we interrogated the Regulome database (Boyle et al. 2012), which provides a means to predict the regulatory potential of SNPs across the genome. In this approach, SNPs are given a score based on their likelihood of being variants with regulatory function (see Materials and Methods for details). Hence, we quantified the enrichment of SNPs with predicted regulatory activity that fall within enhancers in different categories of pleiotropy. We noted that there is a positive relationship between the degree of pleiotropy of enhancers and the level of enrichment in SNPs with predicted regulatory function (Figure 4). When using only SNPs in the most stringent category of Regulome database (category 1a: SNPs that (i) are eQTLs, (ii) match TF motifs, (iii) match DNase footprints, (iv) lie within DNase peaks and (v) lie within TF binding peaks) the positive relationship holds, and the enrichment becomes even larger (Figure S5).

**Figure 4.**
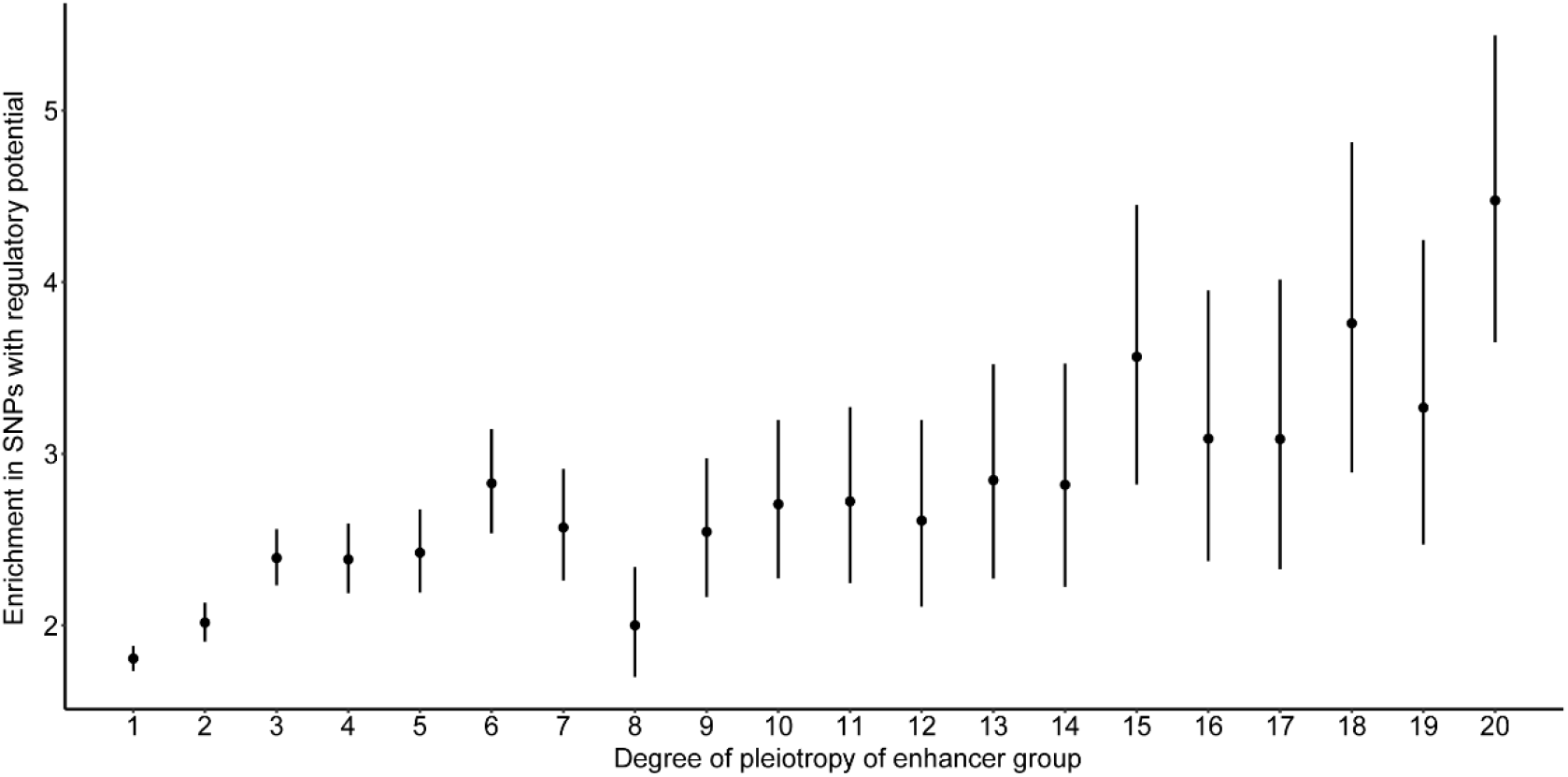
Pleiotropic enhancers are enriched in SNPs with high regulatory potential. The degree of pleiotropy of ENCODE enhancers is positively correlated with the enrichment in SNPs with high regulatory potential (category 1 SNPs of Regulome DB). Black circles represent the odds ratio and bars represent the 95% CI. Enrichments (odds-ratios) were calculated using common SNPs as background (see Materials and Methods for details).

## Discussion

In this study, we predicted active enhancers across human organs by analyzing different types of genomic data. Enhancer predictions were based on either the analysis of eRNA transcription (FANTOM5 enhancers) or the analysis of chromatin architecture (ENCODE enhancers). Remarkably, although these two enhancer sets are different and are derived from different data types, their make-ups are highly similar, which provides a solid basis for our conclusions. Our data suggest that more than 40% of the enhancers in the human genome are pleiotropic. We argue that 40% is a minimum estimation because we mostly considered adult organs (we included just a few fetal biosamples) and because we cannot identify enhancers that are active in a single organ but regulating more than one gene (these context-specific enhancers are pleiotropic as well). However, it is also conceivable that using bulk data from organs imposes a bias in the identification of pleiotropic enhancers. To evaluate this possibility, we calculated the percentage of pleiotropic enhancers using primary cells instead of organs. Notably, the percentage of pleiotropic enhancers calculated with cells is similar, or even larger, than the percentage obtained with organs. Hence, it is unlikely that the estimation of the percentage of pleiotropic enhancers in organs is inflated by the contribution of the same cell types present in multiple organs (e.g., fibroblasts).

A further analysis of the abundance of context-specific versus pleiotropic enhancers per organ uncovered that there is a wide variability in the proportion of context-specific enhancers per organ, and that the majority of active enhancers in a particular organ are pleiotropic. Furthermore, when trying to comprehend the variability in the proportion of context-specific enhancers among organs, we showed that this proportion might be partly explained by the occurrence of context-specific genes in that same organ.

Altogether, the analysis of ENCODE and FANTOM5 enhancers suggests that pleiotropic enhancers are ubiquitous regulatory elements in the human genome. The notion that such a high proportion of human enhancers can be pleiotropic suggests that small regions of regulatory DNA may contain abundant regulatory information and that these regions evolve under important evolutionary constraints, as was previously observed for some mammalian (Fish et al. 2017; Hiller et al. 2012) and human (Radke et al. 2021) pleiotropic enhancers.

Previous studies determined that pleiotropic enhancers have more density and diversity of TF binding motifs (Fish et al. 2017; Singh & Yi 2021). Clearly, this pattern makes sense, since it is logical to expect an increase in the number of TFBSs when more expression patterns are encoded within an enhancer. Investigating the possible regulatory function of SNPs within our enhancer pleiotropy categories, we found that the number of SNPs with regulatory potential increases with the degree of pleiotropy of the enhancer, implying that the more pleiotropic an enhancer is, the more regulatory information that enhancer contains. This finding confirms the suitability of our pleiotropy categories.

Although our data indicate that pleiotropic enhancers regulate gene expression in multiple organs, they do not enlighten the mechanisms underlying pleiotropic function. Pleiotropic enhancers may harbor distinct sets of TFBSs for driving gene expression in different organs. Alternatively, the same TFBSs within pleiotropic enhancers might be needed to regulate gene expression in multiple organs (i.e., pleiotropic enhancers bearing pleiotropic TFBSs). Indeed, a recent study showed that pleiotropic SNPs regulate the expression of *IRX3/IRX5* genes in adipose and brain tissue of humans and mice (Sobreira et al. 2021). Furthermore, a comparative analysis of the genetic architecture of hundreds of human complex traits also provides evidence for the existence of pleiotropic SNPs in enhancers of the human genome (Watanabe et al. 2019). Undoubtedly, this and other issues related to the structure and function of pleiotropic enhancers remain to be studied in detail.

## Materials and Methods

### The FANTOM5 enhancer set

The FANTOM5 enhancer set was downloaded as a binary matrix from https://enhancer.binf.ku.dk/presets. Of all the available biosamples, 139 were mapped to 41 organs (Table S1) based on UBERON ontology terms, as described in Andersson et al. 2014. The FANTOM5 enhancers were identified in each biosample as bidirectional transcribed loci that are expressed significantly above genomic background and that are distal from known TSSs and exons of protein coding genes. These elements were subsequently filtered to remove directionally biased elements (those with a higher level of transcription in one of the DNA strands), which often correspond to TSSs of protein-coding genes. We eliminated from this set of consensus elements those that overlap with regions that are in the ENCODE blacklist of the hg19 assembly (https://github.com/Boyle-Lab/Blacklist). In total, we kept 14814 putative enhancer elements. A subset of these elements has been validated with reporter constructs in HeLa cells (Andersson et al. 2014). Enhancers were considered active in an organ if they were active in at least one of the biosamples that comprise the organ. Enhancers included in the pleiotropy category *n* were those predicted to be active in *n* organs. The size distribution of FANTOM5 enhancers is shown in figure S2.

### Generation of the ENCODE enhancer set and definition of pleiotropic enhancers

For generating the ENCODE enhancer set we used high quality DNase-seq and H3K27ac ChIP-seq data from 37 tissues from the ENCODE Project. We downloaded alignment files in bam format for DNase-seq data (see table S4 for details) and peak files in bed narrowPeak format for H3K27ac ChIP-seq data (see table S5 for details) from https://www.encodeproject.org. For each DNase-seq experiment, we called peaks with MACS2 (Zhang et al. 2008) with --min-size 50 -q 0.01 -f BAMPE for paired-end experiments and --min-size 50 -q 0.01 -f BAM --extsize 200 for single-end experiments. Since the precise location and width of a given accessibility peak may vary between tissues and technical replicates of the same tissue, we developed a method for merging elements from different tissues by searching for overlaps between the summits of accessible elements rather than overlaps between whole elements. To achieve this, we constructed summit confidence intervals for each element in each tissue by using the variability in the position of summits of the same accessible element on the different replicates. Firstly, for each tissue with replicates we estimated the signal-to-noise ratio (SNR) by calculating the proportion of reads inside peaks (analogous to the Fraction of Reads in Peaks score for ATAC-seq) with featureCounts (Liao et al. 2014) using the following parameters: --p --countReadPairs for paired-end experiments and default parameters for single-end experiments. We then intersected the highest SNR experiment MACS2 called peaks with the available H3K27ac peaks for the same tissue (when multiple replicates of H3K27ac ChIP-seq experiments where available, we used only those H3K27ac peaks represented in more than half of the replicates). We then intersected these DNase-H3K27ac peaks with the DNase-seq peaks of the other replicates with bedmap (Neph et al. 2012) using parameters --fraction-either 0.5 and created a gappedPeak file with a custom-made R script that represents the positions of the summits of the same element in the different replicates. DNase-H3K27ac peaks only present in the highest SNR file were not included in subsequent analyses. We called intervals in gappedPeak files ‘summit confidence intervals’. Summits of tissues with only one DNase-seq were only considered if the element overlapped with H3K27ac peaks in that same tissue. By overlapping the summit confidence intervals between tissues, we created ‘summit clusters’ present in multiple tissues. We created consensus elements by merging all the accessible elements (MACS2 peaks) that contribute to each summit cluster. Consensus elements overlapping by more than 80% of the sequence were merged. The 37 ENCODE tissues were mapped to 19 UBERON organs and one cell ontology category (Lymphocyte; CL:0000542) (Table S2). Enhancers active in an organ are consensus elements predicted to be active in at least one of the tissues that comprise that organ. With these data we constructed a binary matrix in which rows represent consensus elements and columns represent organs. Pleiotropic enhancers were defined as consensus elements predicted to be active in 2 or more organs. The code generated to create the consensus elements from DNase-seq and H3K27ac ChIP-seq data can be found at https://github.com/frankel-lab/human_pleiotropic_enhancers. We eliminated from this set of consensus elements those that overlap with regions that are in the ENCODE blacklist of the hg38 assembly (https://github.com/Boyle-Lab/Blacklist), those that are located at less than 500 bp from annotated transcription start sites and those that overlap exons (including putative enhancers that overlap exons does not affect the analyses, data not shown). The final set of putative enhancers consisted of 334189 elements.

### Comparisons between FANTOM5, ENCODE and ABC enhancer sets

The ABC enhancer set was downloaded from https://www.engreitzlab.org/resources. We mapped the biological samples to the corresponding UBERON organs (Table S6) in order to compare with FANTOM5 and ENCODE enhancers. ABC enhancer regions overlapping by more than 80% were merged with bedtools (Quinlan & Hall 2010). In total, we used 176220 ABC elements. FANTOM5 and ABC enhancers were lifted to the hg38 human genome assembly with UCSC liftOver (https://genome.ucsc.edu/cgi-bin/hgLiftOver) for comparing with the coordinates of the ENCODE enhancers. The overlap fraction was calculated by intersecting enhancers active in each organ (comparing two enhancers sets) and then dividing by the size of the smaller set (Table S7).

### Calculation of the percentage of pleiotropic enhancers

To evaluate how the proportion of pleiotropic enhancers changes with the number of organs considered we took all possible combinations of organs from 2 to *n*, where *n* is the total number of organs (20 for the ENCODE enhancers and 41 for the FANTOM5 enhancers) and calculated the proportion of enhancers active in 2 or more organs. For *k* organs, there exists *n* choose *k* combinations of organs. For *k* values with more than 2000 combinations, we estimated the distribution of the proportion of pleiotropic enhancers with 2000 randomly chosen sets of size *k* (without repetition).

We calculated the proportion of pleiotropic enhancers (cCRE-dELS) for 17 cell types (Table S2). The ENCODE cCRE-dELS (n= 290876) in these cells were defined by Moore et al (2020). We explicitly excluded from the analysis data stemming from cancer derived cell lines. We also calculated the proportion of pleiotropic enhancers in 39 cell types (69 primary cell cultures were combined to form 39 different cell types, see table S1)(Andersson et al. 2014). For comparing the degree of pleiotropy of enhancers in cell types and organs we analyzed the 11400 enhancers that were detected as active in both cell types and organs (Andersson et al. 2014).

### Correlation between the proportion of organ-specific enhancers and organ-specific TSSs/transcripts

For FANTOM5 enhancers, we used the same CAGE-seq libraries that were used to identify enhancers to estimate the proportion of context-specific TSSs (reads that correspond to first exons of genes). We used the RefSeq curated gene set (ncbiRefSeqCurated, assembly hg19) from the UCSC site (https://genome.ucsc.edu/). We extended each TSS 100 bp in both directions and merged overlapping elements that were on the same DNA strand with bedtools (Quinlan & Hall 2010) (we obtained similar results without merging, data not shown). We obtained a total of 25797 extended TSSs. For each biosample, we counted CAGE-seq reads overlapping extended TSSs in a strand-specific manner with featureCounts and compared the counts of each TSS with the count distribution of 1477250 random genomic elements of the same size (we excluded exons and TSSs from GRCh38 and all the ENCODE and FANTOM5 enhancers from this random genomic elements) generated with bedtools. We computed an empirical p value for each TSS in each biosample as the fraction of random elements with the same or greater number of reads than the TSS in that biosample. We corrected these empirical p values with the Benjamini-Hochberg procedure and considered that a TSS is active in a biosample if its corrected p value was 0.01 or less (we obtained similar results using thresholds of p<0.005 or p<0.001; data not shown).

For ENCODE enhancers we used ENCODE RNA-seq data available for 25 of the 37 tissues that were used to create the enhancer set (Table S8). We downloaded FPKM normalized counts and only kept genes with protein products annotated by the HUGO Gene Nomenclature Committee (19167 genes). When replicates of the same tissues were available, we used the median FPKM for each gene. In order to use a uniform threshold across tissues to decide when a gene is expressed or not, we used the zFPKM R package (Hart et al. 2013) to normalize FPKM. Genes were considered to be expressed if their zFPKM was above -2 (we obtained similar results using -3.5, -3, -2.5, -1,0 and 1 as thresholds; data not shown). We regard a gene as context-specific when the gene is expressed in a single organ (for both the FANTOM5 and ENCODE analyses).

### RegulomeDB SNP analysis

SNPs with regulatory potential were downloaded from the RegulomeDB site (https://regulomedb.org/files/TSTFF344324). We lifted the hg19 positions to the hg38 assembly using UCSC liftOver. For this analysis we considered SNPs in category 1 (SNPs that are eQTLs and that have at least one additional source of evidence for being regulatory) and the most stringent category of RegulomeDB, which is category 1a (Boyle et al. 2012). Odds ratios were calculated for each pleiotropy category as the RegulomeDB SNPs contained in the category divided by the non-RegulomeDB SNPs (1000 Genomes SNPs with a MAF > 0.05 in Europeans) in that same category divided by the same ratio for the rest of the genome. In total we considered 71263 SNPs in category 1 and 978 SNPs in category 1a, and 7894464 common variants. All SNP count operations were performed with bedmap.

## Acknowledgments

We thank Justin Crocker, Hernan Dopazo and members of the Frankel lab for helpful comments on the manuscript. This work was supported by grants PICT 2016-0414 and PICT 2019-00497 (ANPCyT, Argentina) to NF.

## Competing interests

The authors have no competing interests to declare

## Data availability statement

The data underlying this article are available in the article and in its online supplementary material.

## Notes

### Competing Interest Statement

The authors have declared no competing interest.

