## Supplementary figures for "Pleiotropic enhancers are ubiquitous regulatory elements in the human genome"

Figure S1

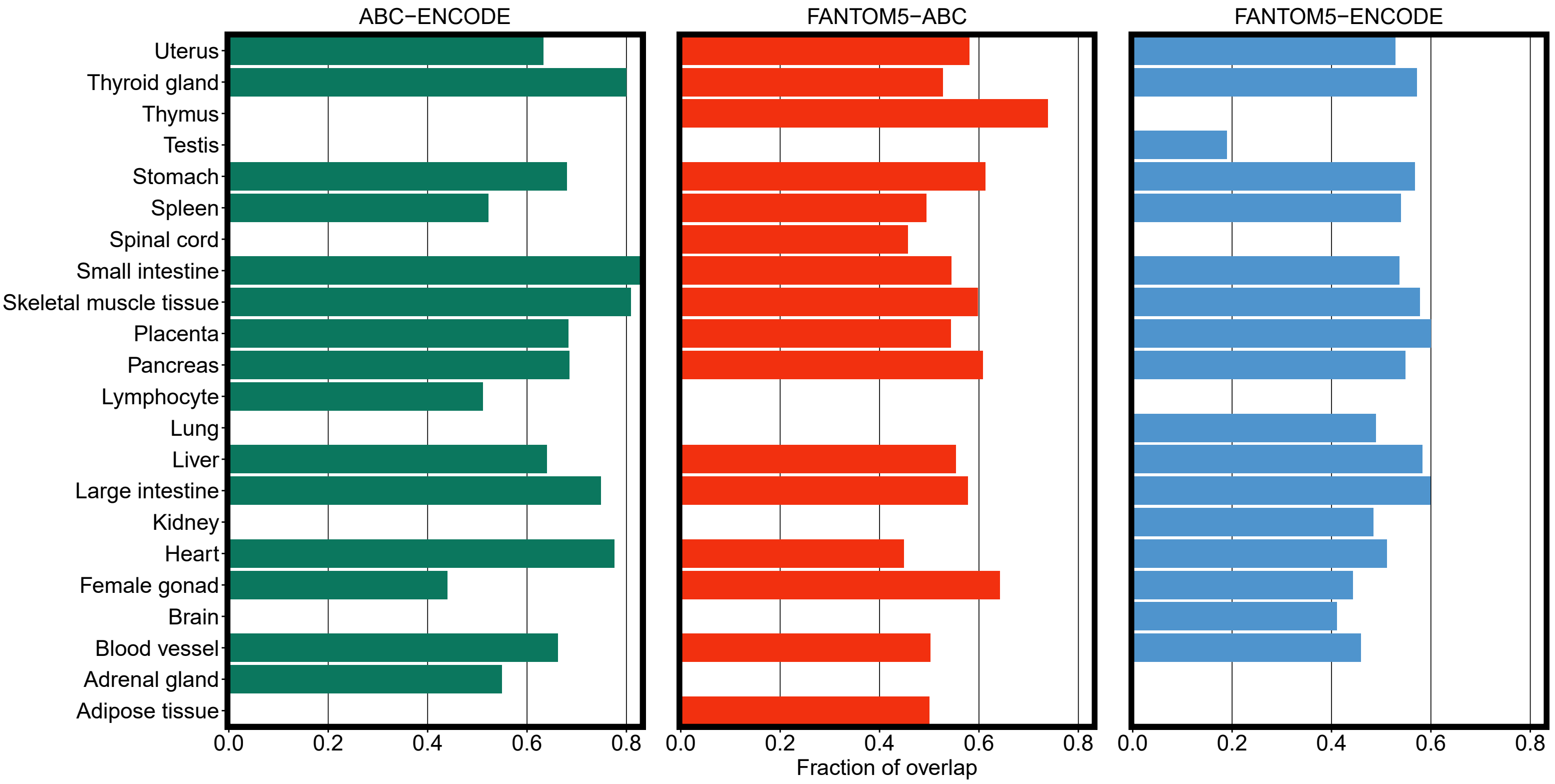


Figure S1. Overlap between enhancer sets. For each organ, we calculated the fraction of overlap as the number of overlapping elements on the smaller set divided by the size of the smaller set.

Figure S2

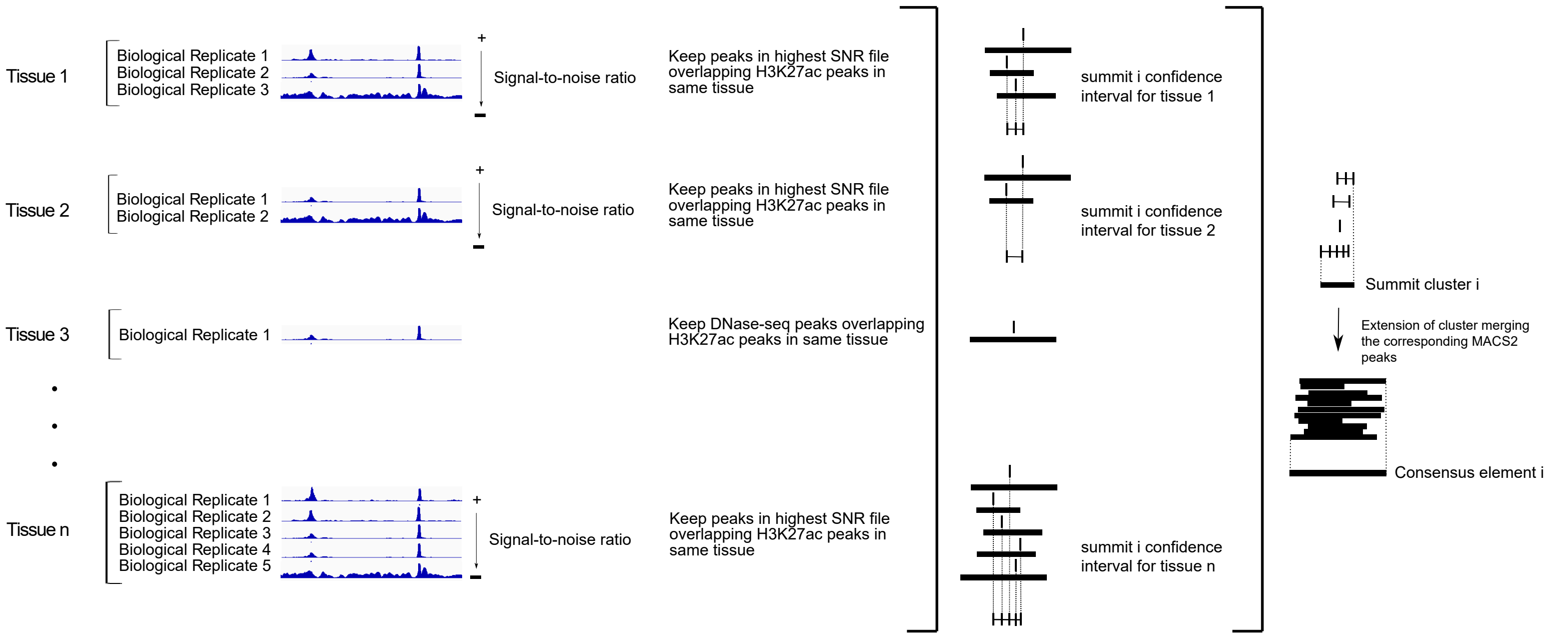


Figure S2. Pipeline for delimiting consensus elements. The diagram illustrates the steps for delimiting the sequence of pleiotropic enhancers, considering all organs in which an enhancer is active.

Figure S3

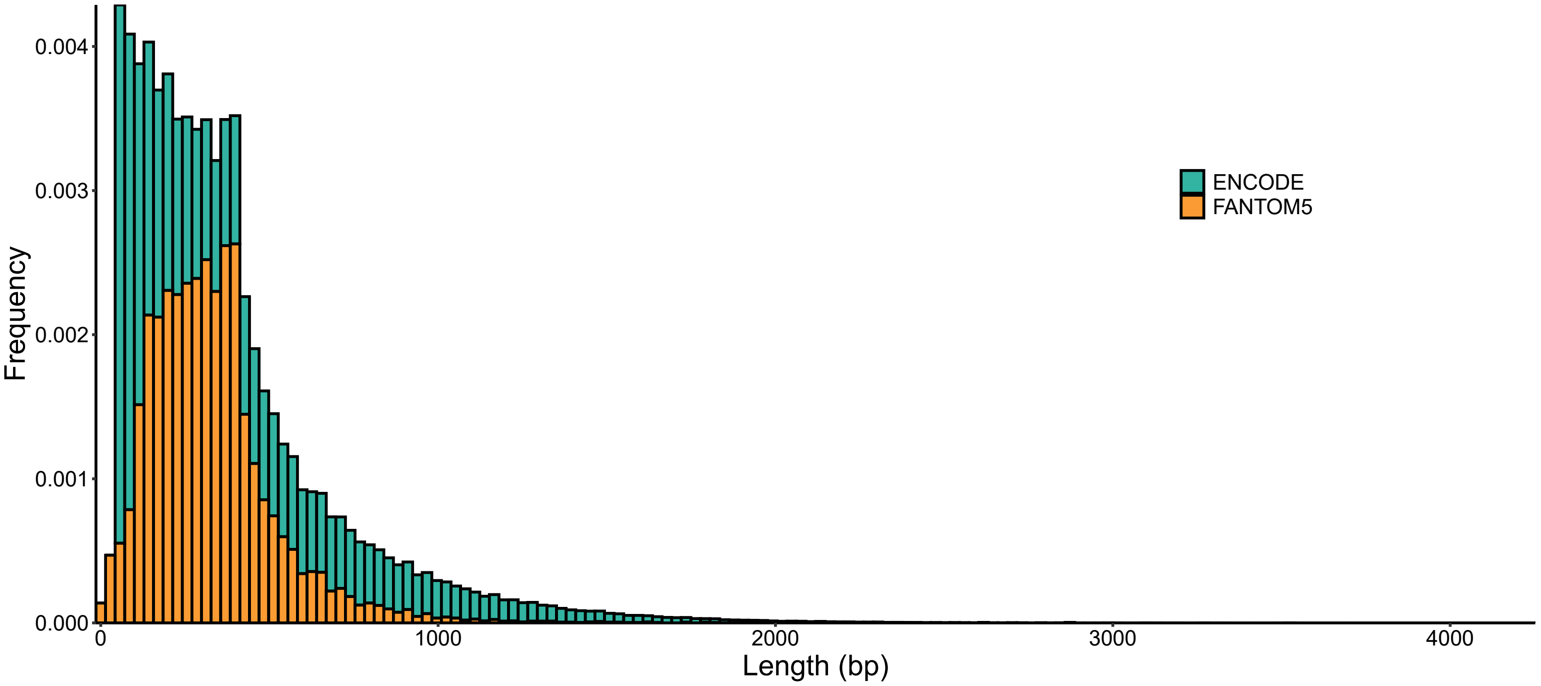


Figure S3. Size distribution of predicted enhancers. Sizes in base-pairs for ENCODE consensus enhancers (green) and FANTOM5 (orange) enhancers.

Figure S4

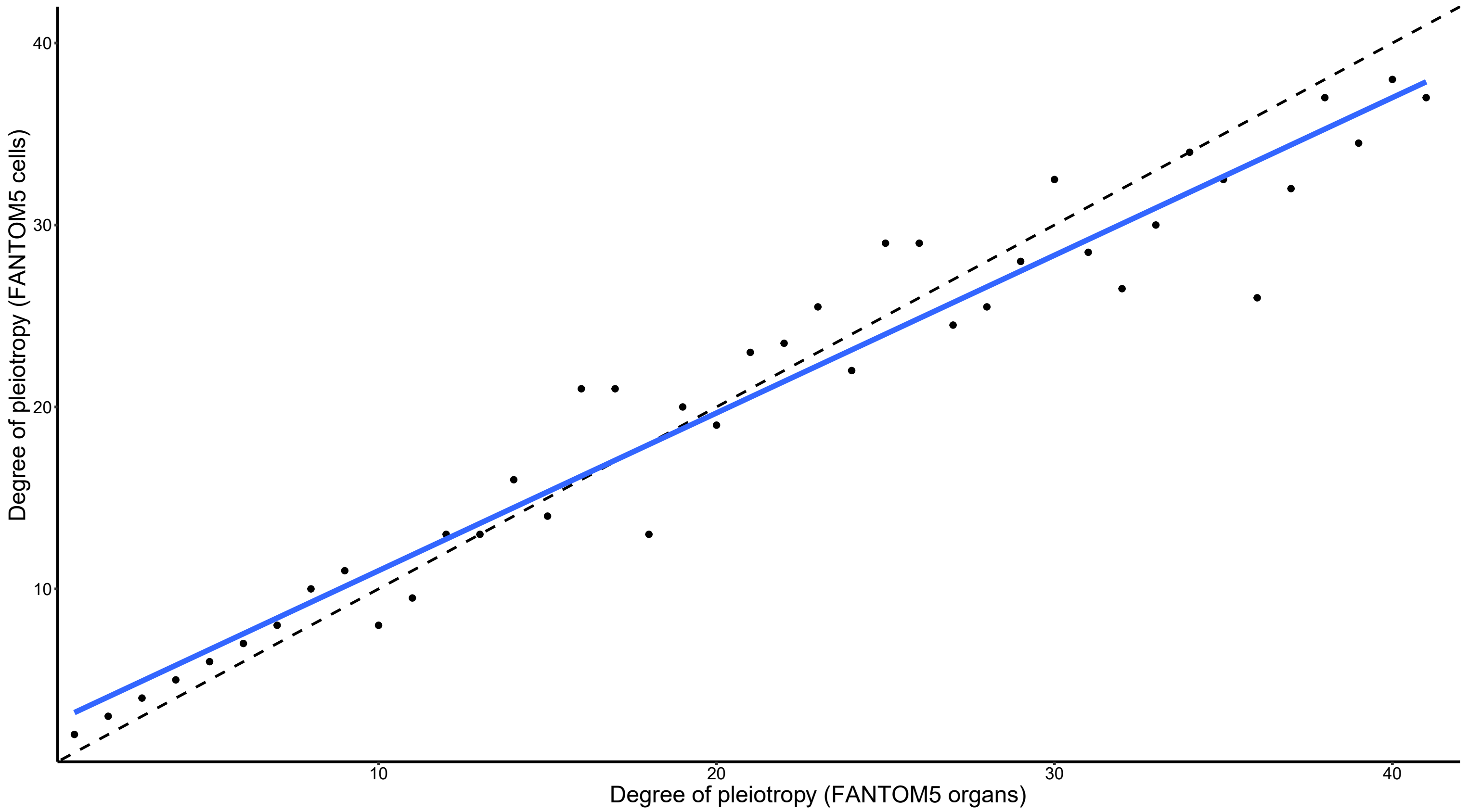


Figure S4. Comparison of the estimated degree of pleiotropy for FANTOM5 enhancers calculated with data from organs (X axis) and data from cell types (Y axis). The black dashed line marks the diagonal, which is the expectation if organ data and cell types data provided exactly the same estimate. The blue line represents the best fit for the median values per category (black dots).

Figure S5

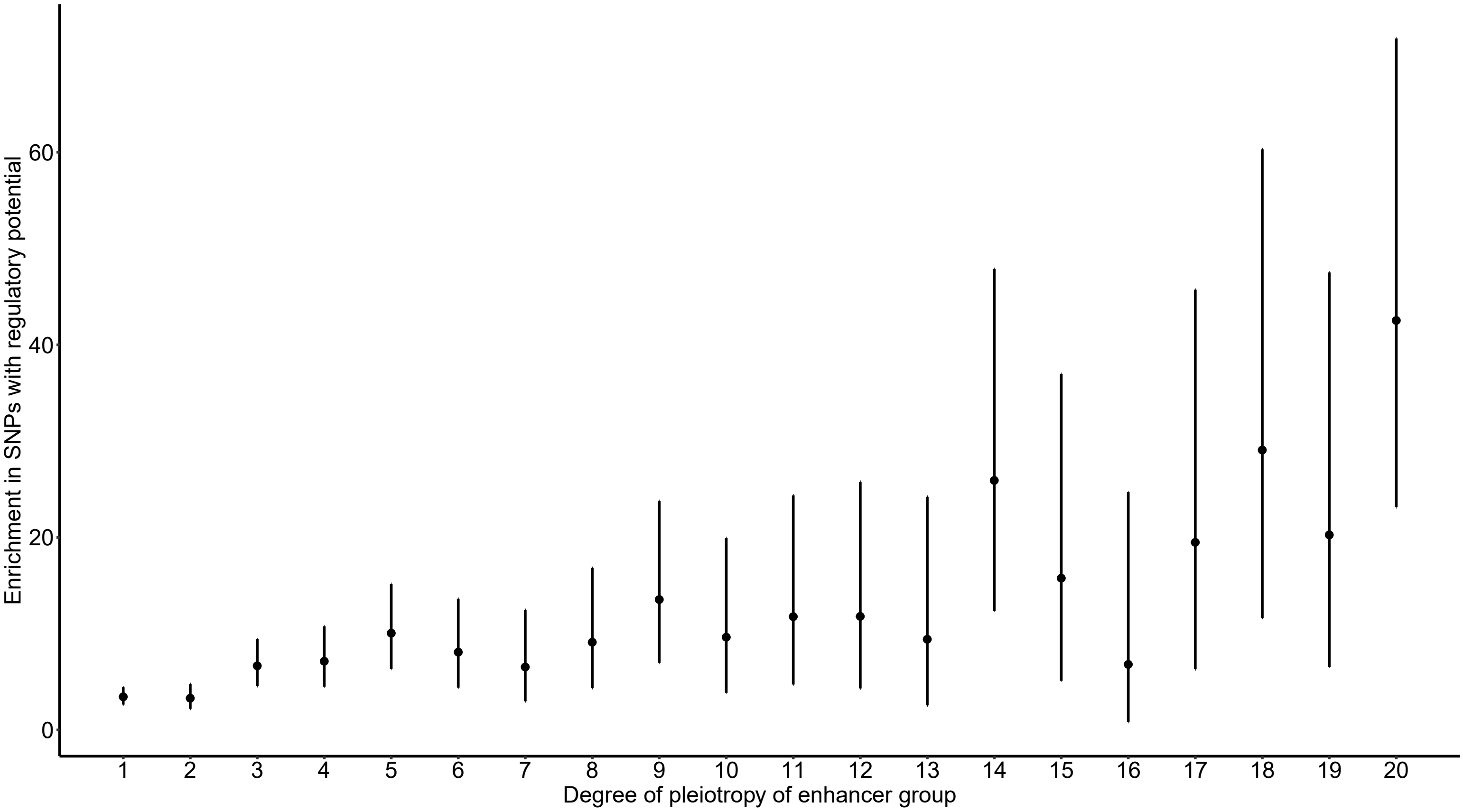


Figure S5. Pleiotropic enhancers are enriched in SNPs with high regulatory potential. The degree of pleiotropy of ENCODE enhancers is positively correlated with the enrichment in category 1a SNPs of regulome DB. Black circles represent the odds ratio and bars represent the 95% CI.
